# MicroRNA regulation of stress-survival signalling and protein quality control in human heatstroke

**DOI:** 10.64898/2026.06.25.734416

**Authors:** Maria Gomez, Saeed Al Mahri, Mashan L. Abdullah, Shuja Shafi Malik, Marwa Abdelhakim, Saber Yezli, Robert Hoehndorf, Abderrezak Bouchama

## Abstract

Heatstroke is a life-threatening condition in which heat-shock and unfolded-protein responses are strongly activated but fail to prevent proteostasis disruption and severe cellular injury. Whether post-transcriptional regulation contributes to this mismatch remains unknown. We integrated small RNA sequencing with mRNA profiling in peripheral blood mononuclear cells from patients with classical heatstroke and matched heat-exposed controls recruited during the Hajj pilgrimage. mRNA profiling was performed in 19 cases and 19 controls, and miRNA sequencing in 17 cases and 16 controls from the same cohort. Differentially expressed miRNAs were integrated with 4,462 differentially expressed mRNAs using high-confidence inverse-expression miRNA-mRNA pairs. Twenty-six miRNAs mapped to 376 mRNA targets, forming 414 regulatory pairs and two opposing programmes. Programme A, comprising 16 downregulated miRNAs, was associated with activation of PI3K-mTOR, NRF2 oxidative stress and HIF-1α signalling, consistent with stress-survival signalling. Programme B, comprising 10 upregulated miRNAs, was associated with suppression of stress-granule components and fatty-acid β-oxidation genes, consistent with impaired protein quality control and reduced metabolic flexibility. miR-92a-3p emerged as a central regulatory node, and its target PIK3R3 connected 9 of the 10 enriched pathways. These findings suggest a post-transcriptional regulatory layer that could contribute to the limited protection afforded by activated stress defences in human heatstroke.

**Funding:** King Abdullah International Medical Research Center (RC18/373/R).

**Key points:**

- Heatstroke can occur even when major cellular stress defences are activated, raising the question of why heat-shock and unfolded-protein responses are insufficient to prevent injury under extreme heat.
- To understand this mismatch, we extend our previous human heatstroke transcriptomic study by analysing post-transcriptional regulation by microRNAs (miRNAs).
- The miRNA-regulated network showed activation of stress-survival signalling pathways, including PI3K–mTOR, NRF2 and HIF-1α, while stress-granule and fatty-acid β-oxidation pathways were attenuated.
- Translation arrest was active but stress-granule signalling was suppressed. Normally, translation arrest generates pools of non-translating mRNAs that stress granules sort, store or return to translation when conditions improve; their uncoupling therefore suggests impaired coordination of the cellular stress response.
- These findings suggest a post-transcriptional trade-off in human heatstroke, in which short-term stress-survival signalling is favoured at the cost of protein quality control, metabolic flexibility and recovery capacity under extreme heat.

## Introduction

Heatstroke is a life-threatening condition that results when heat gain exceeds the capacity for heat dissipation, either through passive exposure to extreme environmental heat in classic heatstroke or through excessive endogenous heat production during strenuous exercise in exertional heatstroke.(Bouchama *et al*., 2022) Alongside thermoregulation, survival during extreme heat depends on cellular stress-response systems that preserve protein homeostasis and energy metabolism under thermal load.(Richter *et al*., 2010; Bouchama *et al*., 2022)

Central to this cellular defence is the heat-shock response (HSR), a conserved cytoprotective programme in which molecular chaperones limit protein misfolding, support refolding or degradation of damaged proteins, and maintain proteostasis under thermal stress.(Lindquist & Craig, 1988; Richter *et al*., 2010; Mahat *et al*., 2016) Heat stress also activates the integrated stress response (ISR) and unfolded-protein response (UPR), which reduce global protein translation, limit the burden of newly synthesized proteins, promote selective stress-adaptive gene expression, and support mRNA triage through stress-granule formation (Walter & Ron, 2011; Costa-Mattioli & Walter, 2020)

We previously reported the peripheral blood mononuclear cell (PBMC) transcriptomic signature of acute human heatstroke. (Bouchama *et al*., 2023) That analysis showed broad differential expression of heat-shock protein, co-chaperone and chaperonin genes, indicating a robust HSR, together with gene-expression changes related to the UPR, DNA repair, energy metabolism, oxidative stress and immunity. (Bouchama *et al*., 2023) These defence signals were observed in patients who had already developed life-threatening heatstroke, with core temperature above 40°C, severe neurological dysfunction and biochemical evidence of multiorgan stress. This clinical-molecular contrast suggests that conserved stress defences were activated but insufficient to prevent severe heat injury. Several mechanisms may contribute to this insufficiency, including persistent proteotoxic stress, impaired clearance of damaged proteins, mitochondrial energy failure, and altered post-transcriptional or translational regulation. Among these mechanisms, microRNAs (miRNAs) are of particular interest because they regulate gene expression post-transcriptionally.

MicroRNAs are endogenous small non-coding RNAs (∼22 nucleotides) that regulate gene expression post-transcriptionally, with individual miRNAs targeting hundreds of mRNAs across multiple pathways simultaneously.(Bartel, 2004; Emde & Hornstein, 2014; Ha & Kim, 2014) miRNAs modulate translation arrest, autophagy, apoptosis, and inflammatory signaling during stress.(Emde & Hornstein, 2014; Si *et al*., 2019) In heatstroke patients, exosomal miRNA profiling identified signatures enriched in inflammation and coagulation pathways.(Li *et al*., 2021) In a rat heatstroke model, integrated miRNA and transcriptomic analysis revealed that differentially expressed miRNAs target pathways governing energy metabolism, the UPR, and *PI3K-Akt* signaling, with the greatest perturbation in animals developing organ injury.(Permenter *et al*., 2019) Integrated miRNA–mRNA networks have also been reconstructed in rat small intestine after heat injury(Yu *et al*., 2011) and in non-lethal heat-stressed shrimp, (Boonchuen *et al*., 2020) and wheat.(Mishra *et al*., 2023) However, no study has yet integrated miRNA and mRNA expression into a unified regulatory network in human heatstroke, nor examined how such networks might organize clinically relevant stress pathways in patients.

Leveraging transcriptomic data from our previously characterised cohort of heatstroke patients and heat-exposed controls (Bouchama *et al*., 2023), we integrated newly generated miRNA profiles to examine whether miRNA-mediated regulation may clarify why a robust heat-shock and unfolded-protein response is insufficient to prevent severe heat injury. Specifically, we asked whether differentially expressed miRNAs are linked to mRNA targets within cellular stress-response, proteostasis, and metabolic pathways. To address this question, we performed integrated miRNA-mRNA network analysis in PBMCs from heatstroke patients and matched heat-exposed controls recruited during the Hajj pilgrimage. (Bouchama *et al*., 2023)

## Methods

### Study design and participants

This study extends our previously published research on the clinical and transcriptomic signatures of heatstroke.(Bouchama *et al*., 2023) Clinical characteristics, biochemical parameters, and mRNA expression profiles have been reported in detail;(Bouchama *et al*., 2023) here we present miRNA expression profiles and integrated miRNA–mRNA regulatory network analysis from the same cohort.

The study was conducted at Mina Emergency Hospital in Mecca, Saudi Arabia, during the August 2017 pilgrimage season. The Institutional Review Board of King Abdullah International Medical Research Centre approved the protocol (IRBC/817/17), and all participants provided written informed consent in accordance with the Declaration of Helsinki.

### Heatstroke patients

Nineteen adult patients with heatstroke were enrolled based on rectal temperature >40.1°C on admission, neurological dysfunction (delirium, convulsions, or coma), and documented environmental heat exposure. Patients with cardiac arrest or who declined consent were excluded.

### Control subjects

Nineteen non-heatstroke controls were recruited from pilgrims who were friends or relatives of heatstroke patients and were matched for environmental exposure. All participants resided within the same geographic area, wore similar clothing, consumed comparable diets, and engaged in similar pilgrimage activities.

### Sample size and blinding

No formal a priori power calculation was performed. The sample size was determined by the availability of eligible heatstroke cases and matched controls collected under standardized field conditions. Cohort size is consistent with prior human transcriptomic studies of heat stress (Bouchama *et al*., 2017) and exertional heat injury.(Sonna *et al*., 2004)

This was an observational case–control study; randomisation was not applicable. Laboratory processing followed standardized protocols for all samples. RNA extraction, microarray hybridisation, and sequencing library preparation were performed using identical procedures irrespective of group assignment. Uniform preprocessing and quality-control procedures were applied across samples, and multiple testing was controlled using the false-discovery rate (FDR).

### Sample collection and processing

Blood samples were collected from heatstroke patients upon arrival at the cooling unit before cooling treatment (T0) and again within four hours after cooling (T1), as previously described. (Bouchama *et al*., 2023) The present miRNA analysis was restricted to T0 samples because available funding supported small RNA sequencing only at the admission time point. Control samples were collected upon enrolment. PBMCs were isolated using Leucosep tubes (Greiner Bio-One) according to the manufacturer’s instructions. Total RNA, including small RNAs, was extracted using TRIzol Reagent (Thermo Fisher Scientific) followed by purification with the SV Total RNA Isolation System (Promega). RNA integrity was assessed using the Agilent 2100 Bioanalyzer; samples with RNA integrity number (RIN) >7 were used for downstream analysis.

### mRNA expression profiling

Gene expression profiling was performed using the Human Clariom D Array (Thermo Fisher Scientific) as previously described.(Bouchama *et al*., 2023) For the present study, raw CEL files were reprocessed using Brainarray custom chip definition files (ClariomDHuman_Hs_REFSEQ, version 25·0·0) with background correction and RMA normalisation using the Affy and Affycoretools R packages. Genes with low expression intensity and low variance were excluded to reduce uninformative probes. Sex and age were included as covariates in the limma linear models, as both influence the heatstroke transcriptomic response in this cohort, with distinct age-related and sex-specific differential expression patterns. (Bouchama *et al*., 2025; Gomez *et al*., 2025). Laboratory preprocessing batch was also included as a covariate. Differential expression was assessed with empirical Bayes moderation and Benjamini–Hochberg correction, yielding 4,462 differentially expressed mRNAs (FDR < 0·05). This represents a more conservative set than the 8,854 genes identified under the initial analytical framework using Brainarray version 22 without covariate adjustment.(Bouchama *et al*., 2023) The reduction reflects updated probe-to-gene annotation and explicit modelling of technical and demographic covariates. Core pathway enrichments including activation of the HSR, UPR, and suppression of oxidative phosphorylation were concordant across analytical approaches. The batch-adjusted expression data are publicly available in GEO (GSE317880), and all miRNA–mRNA integration analyses reported here use this dataset.

### miRNA sequencing and analysis

Small RNA libraries were prepared using the QIAseq miRNA Library Kit. Of the 38 participants with mRNA data, 33 (17 heatstroke, 16 controls) yielded miRNA libraries meeting quality criteria; 5 samples were excluded owing to library preparation failure. Sequencing was distributed across two Illumina platforms: 8 samples (4 heatstroke, 4 controls) on MiSeq (single-end, 50 bp) and 25 samples (13 heatstroke, 12 controls) on NovaSeq 6000-SP (single-end, 50 bp). Sequencing platform was included as a covariate in the differential expression model. Raw reads were processed using Cutadapt (v4.3) for adapter removal and quality trimming, aligned to the human genome (GRCh38/hg38) using Bowtie 2 (v2.5.1), and quantified using miRDeep2 (v0.1.3). Top-expressed hairpins were identified following TMM normalisation. Differential expression was assessed using DESeq2 (v1.12.3) in R (v4.2.2), which applies size-factor normalisation based on a negative binomial distribution model. miRNAs with FDR-adjusted P < 0·05 and |logCFC| ≥ 1 were considered significantly differentially expressed. The miRNA expression data were deposited in GEO (GSE324277).

### miRNA–mRNA regulatory network construction

Ingenuity Pathway Analysis (IPA, QIAGEN, version 2026-02, content version 2025Q4) was used throughout. From the initial upload, 42 miRNAs mapped to the IPA knowledge base. After applying expression cutoffs, 28 miRNAs were analysis-ready, of which 26 (10 upregulated, 16 downregulated) had targeting information (Table S1). The miRNA Target Filter was configured with the 4,462 DE mRNAs. Target predictions were restricted to Tarbase and TargetScan Human with confidence levels of “Experimentally Observed” or “High (predicted).” Only inverse expression pairings were retained. This sequential filtering (Figure 1B) yielded 414 high-confidence miRNA–mRNA pairs involving 376 unique mRNA targets. To support reproducibility, the complete list of the 414 miRNA–mRNA pairs and the Ingenuity Pathway Analysis target-filter configuration are provided as Supplementary Table S2. Cross-referencing against miRTarBase, a curated database of experimentally validated miRNA–target interactions, was used to identify pairs with independent experimental support, including reporter-assay evidence where available.(Cui *et al*., 2025) IPA groups miRNAs sharing seed sequences (nucleotides 2–8); consequently, targets listed under hsa-miR-363-3p are functionally attributable to the miR-92a-3p seed family (seed: AUUGCAC).

**Figure 1.**
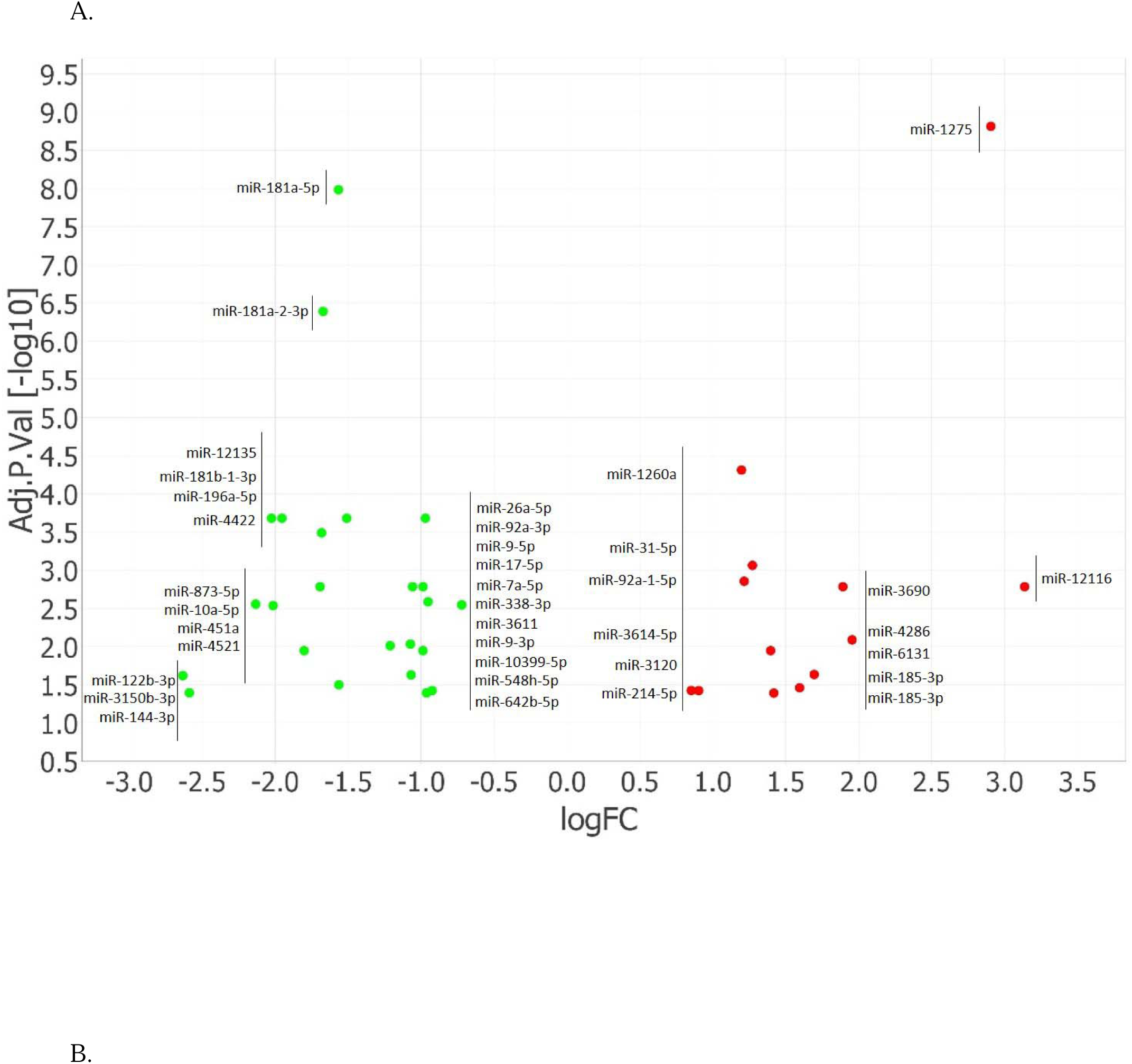
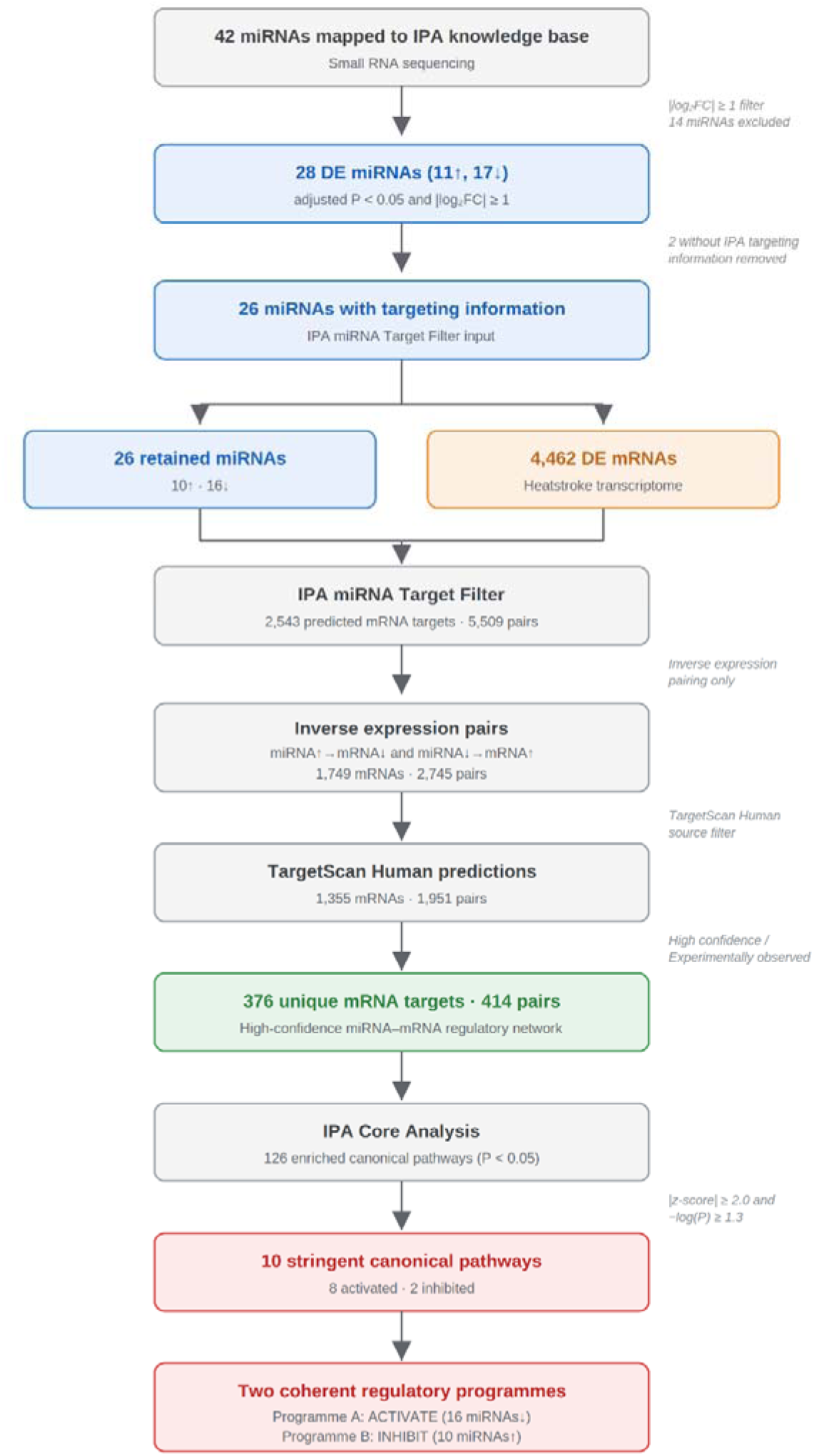
miRNA expression and regulatory network construction in heatstroke. Volcano plot of miRNA expression in PBMCs from heatstroke patients (n = 17) and non-heatstroke controls (n = 16) at admission before cooling (T0). The x-axis shows logFC and the y-axis shows −log10 adjusted P value. Red nodes indicate upregulated miRNAs and green nodes indicate downregulated miRNAs. Forty-two miRNA identifiers mapped to the IPA knowledge base; six were flagged by IPA as duplicate/grouped mappings, in which more than one dataset identifier maps to the same molecular node in the Global Molecular Network. Accordingly, the volcano plot displays 37 visually distinct nodes. (B) Sequential filtering pipeline for miRNA–mRNA regulatory network construction. From the 42 mapped miRNA identifiers, 28 met the predefined differential-expression criteria of adjusted P < 0.05 and |logFC| ≥ 1. Two lacked targeting information in IPA, leaving 26 miRNAs for downstream integration with 4,462 differentially expressed mRNAs. These 26 miRNAs comprised 10 upregulated and 16 downregulated miRNAs. The final high-confidence inverse-expression network contained 376 unique mRNA targets forming 414 miRNA–mRNA pairs and yielded 10 canonical pathways meeting stringent dual criteria (|z-score| ≥ 2.0 and −log(P) ≥ 1.3).

### Canonical pathway analysis

The 376 miRNA-regulated targets were subjected to IPA Core Analysis. Pathway enrichment was assessed using Fisher’s exact test (−log(P) ≥ 1.3). Pathway directionality was predicted using the IPA z-score algorithm; pathways with z-score ≥ +2.0 were considered activated and z-score ≤ −2.0 inhibited. The same Core Analysis was performed on the full heatstroke transcriptome (4,462 DE mRNAs) for comparison.

### miRNA–pathway attribution

Individual miRNA networks were generated in IPA and overlaid with the stringent canonical pathways. Target genes were displayed as nodes with pathway membership indicated by overlaid canonical pathway annotations. Edge color denoted regulatory direction: blue for upregulated miRNAs repressing targets, orange for downregulated miRNAs with de-repressed targets.

### Statistical analysis

Clinical and biochemical variables were summarised as mean ± SD or median (IQR), as appropriate. Between-group comparisons were performed using the Student’s t test, Chi-square test, or Wilcoxon rank-sum test, as appropriate, with two-sided P <0.05 considered statistically significant. These analyses were conducted in SAS version 9.4 (SAS Institute, Cary, NC, USA), as previously reported.

For miRNA differential expression, FDR was controlled at 5% using the Benjamini-Hochberg procedure. Pathway enrichment P values were calculated using Fisher’s exact test with right-tailed testing. Network visualizations were exported at 600 dpi.

## Results

### Participant characteristics

Demographic and clinical characteristics of heatstroke patients (n=19) before cooling (T0) and non-heatstroke controls (n=19) have been reported previously and are summarised in Table 1. miRNA sequencing data were available for 17 heatstroke patients and 16 controls from this cohort. At admission, heatstroke patients presented with severe hyperthermia, elevated hepatic transaminases, renal impairment, and raised muscle tissue-damage markers, consistent with established multiorgan stress.

**Table 1.** Clinical characteristics of heatstroke patients before cooling and non-heatstroke controls.

| Parameters | NHS Controls (N = 19) | HS Patients at T0 (N = 19) | P value |
| --- | --- | --- | --- |
| Age (years) | 50 (41, 55) | 58 (45, 66) | 0.050 |
| Gender (Female/Male) | 8 / 11 | 11 / 8 | 0.330 |
| Core Temperature (°C) | 36.3 ± 0.8 | 41.7 ± 0.8 | <0.0001 |
| Albumin (g/L) | 40 (39, 42) | 40 (39, 42) | 0.45 |
| Alanine Transaminase (U/L) | 15 (13, 26) | 28 (21, 44) | 0.03 |
| Aspartate Transaminase (U/L) | 20 (18, 24) | 40 (25, 67) | 0.001 |
| Bilirubin (µmol/L) | 13.1 (7.4, 14.7) | 13.7 (8.6, 17.4) | 0.37 |
| Creatine Kinase (U/L) | 208 (135, 291) | 326 (135, 874) | 0.11 |
| Creatinine (µmol/L) | 73 ± 13.4 | 133 ± 55.9 | 0.0002 |
| Lactate Dehydrogenase (U/L) | 228 (203, 268) | 337 (272, 390) | 0.0004 |

*Data are presented as median (Q1, Q3) or mean ± SD. HS: Heatstroke before cooling treatment; NHS: Non-heatstroke controls. Lactate Dehydrogenase (LDH) values were missing for 2 heatstroke patients; percentages and summary statistics for LDH were calculated from available data. All other variables were complete.*

### miRNA expression profiling reveals 28 DE miRNAs

We identified 42 miRNAs by small RNA sequencing that mapped to the IPA knowledge base (Figure 1A). After stringent filtering (adjusted P < 0·05, |logCFC| ≥ 1), 28 were differentially expressed between heatstroke patients and controls (11 upregulated, 17 downregulated). The most statistically significant were miR-1275, miR-181a-5p, and miR-181a-2-3p (Figure 1A). The miR-181 cluster (miR-181a-5p, miR-181a-2-3p, miR-181b-1-3p) was uniformly downregulated. Two of the 28 lacked targeting information in the IPA database, leaving 26 miRNAs (10 upregulated, 16 downregulated) for downstream network analysis (Table S1).

### miRNA–mRNA target identification yields 414 pairs

We then integrated the 26 target-annotated miRNAs with the heatstroke transcriptome (4,462 differentially expressed mRNAs) using the IPA miRNA Target Filter. The primary network retained high-confidence inverse-expression pairs: 10 upregulated miRNAs targeting 210 downregulated mRNAs, and 16 downregulated miRNAs targeting 166 upregulated mRNAs, yielding 376 unique mRNA targets connected by 414 miRNA–mRNA pairs (Figure 1B). Thirty-four mRNAs were targeted by two or more miRNAs, indicating convergent regulation.

### Canonical pathway enrichment in the miRNA-regulated subset

We subjected the 376 miRNA-regulated mRNA targets to IPA Core Analysis, which revealed 126 enriched canonical pathways (P < 0·05). Applying stringent dual criteria (|z-score| ≥ 2·0 and −log(P) ≥ 1·3) identified 10 pathways with high-confidence directional predictions (Figure 2A): eight activated and two inhibited.

**Figure 2.**
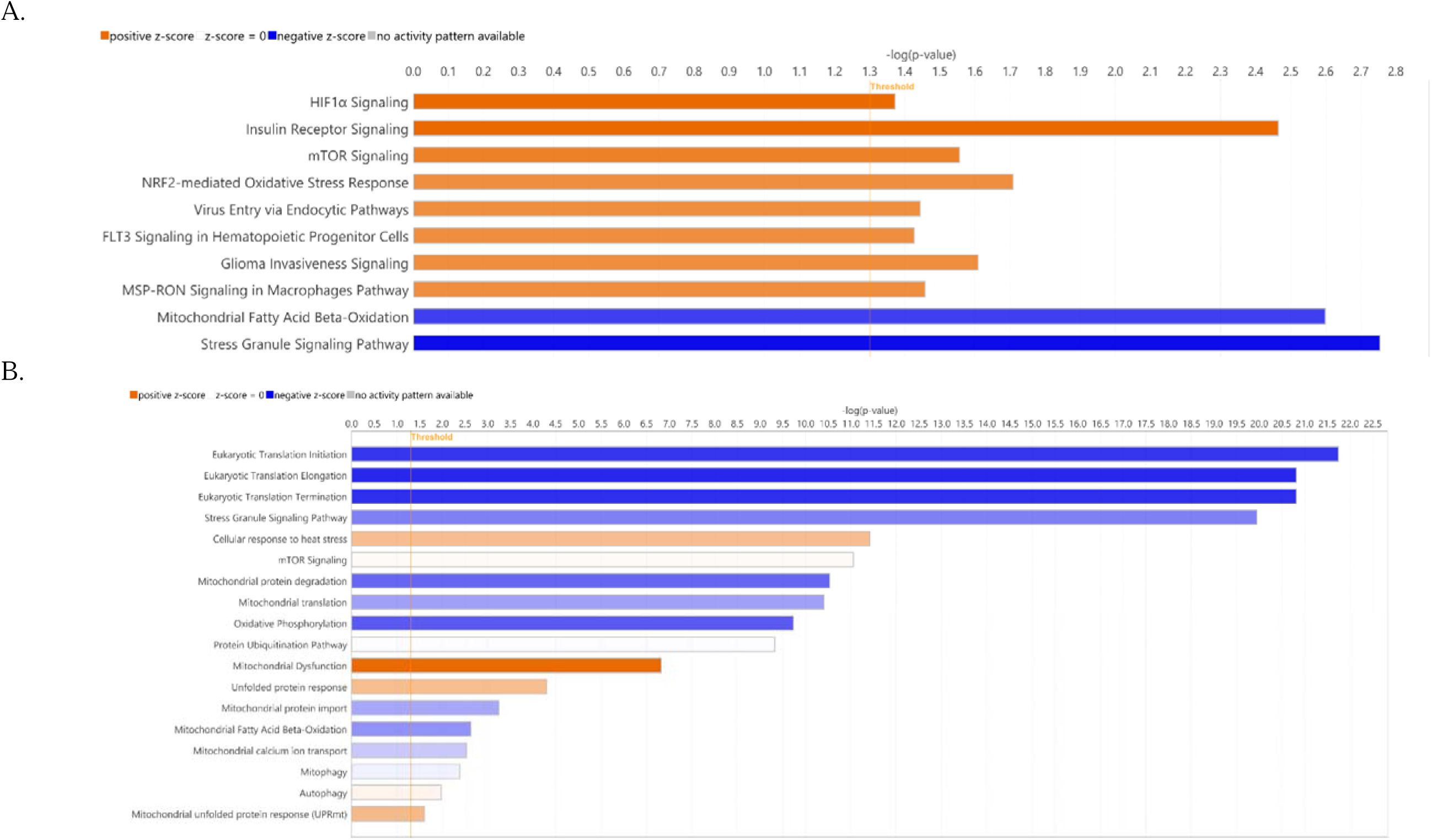
Canonical pathway enrichment in the miRNA network and full transcriptome. (A) Ten canonical pathways meeting stringent criteria from Core Analysis of 376 miRNA-regulated mRNA targets. Orange: predicted activation; blue: predicted inhibition. (B) Selected canonical pathways from Core Analysis of 4,462 DE mRNAs. Gray bars: no significant directional prediction. The transcriptome reveals activated heat-shock/UPR programmes, global translational suppression, and comprehensive mitochondrial dysfuncion. mTOR and clearance pathways were enriched but directionless.

### Comparison with the full heatstroke transcriptome

To contextualise these findings, we compared miRNA-regulated pathway states with the full heatstroke transcriptome (4,462 DE mRNAs) (Figure 2B). The transcriptome showed massive translational suppression (initiation, elongation, and termination all z < −6·8, −log(P) > 20), comprehensive mitochondrial dysfunction, and activated heat-shock/UPR programmes (z = +2·5). Ribosome biogenesis and rRNA processing were strongly inhibited. *mTOR* signalling was highly enriched (−log(P) = 11·1) yet functionally directionless (z = +0·19), and clearance pathways ubiquitination (z = −0·12), autophagy (z = +0·42), mitophagy (z = −0·54) were similarly conflicted. Within the miRNA-regulated subset, *mTOR* met activation criteria (z = +2·24), NRF2 crossed the activation threshold (transcriptome z = +1·96 → miRNA z = +2·00), and suppression of Stress Granule and fatty acid β-oxidation was reinforced.

### miR-92a-3p de-represses phosphoinositide-3-kinase regulatory subunit 3 *(PIK3R3) across 9 of 10 pathways*

Using network overlay, we identified miR-92a-3p as a central node. This downregulated miRNA was linked to 31 targets, 15 of which had independent support in miRTarBase, including Myosin Light Chain Interacting Protein (MYLIP) and dual specificity phosphatase 10 (DUSP10) by reporter assay. In IPA, it is designated hsa-miR-363-3p because of the shared seed sequence AUUGCAC. miR-92a-3p was associated with de-repression of PIK3R3, which participated in 9 of the 10 stringent pathways, all except mitochondrial fatty acid β-oxidation (Figure 3).

**Figure 3.**
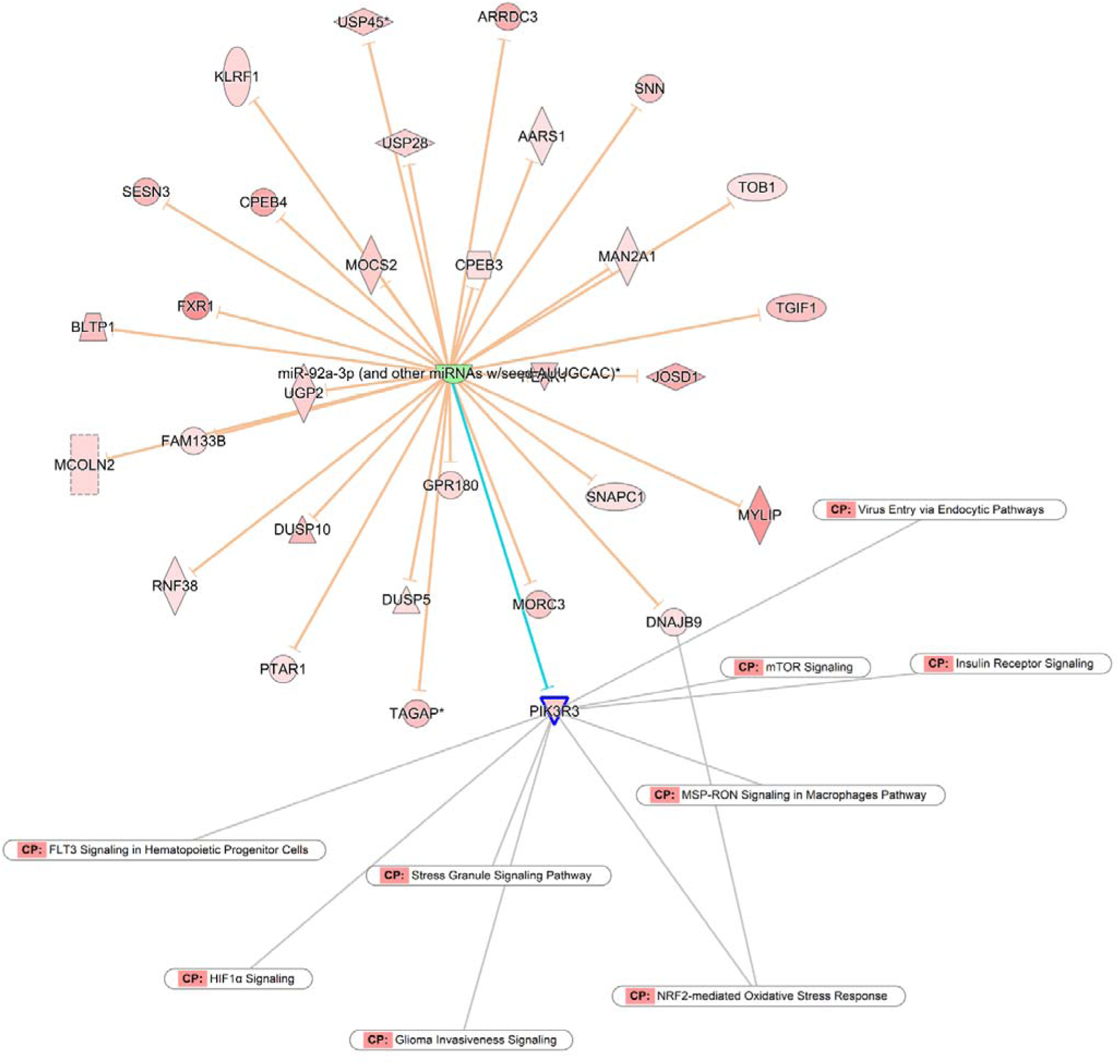
miR-92a-3p regulatory network and PIK3R3 hub architecture. Network overlay of miR-92a-3p (downregulated) and its 31 de-repressed mRNA targets. PIK3R3 connects to 9 of 10 stringent canonical pathways. The sole pathway not reached by PIK3R3 is Mitochondrial Fatty Acid β-Oxidation.

PIK3R3 encodes the p55γ regulatory subunit of class IA PI3K, positioning it at the convergence of mTOR, HIF1α, Insulin Receptor, NRF2, FLT3, Stress Granule, MSP-RON, Virus Entry, and Glioma Invasiveness signalling pathways. PIK3R3’s high connectivity might partly reflect the shared dependence of multiple canonical pathways on PI3K signalling within the IPA knowledge base. Beyond PIK3R3, four additional genes each connected to 3–7 stringent pathways: eukaryotic translation initiation factor 4E (EIF4E), de-repressed by miR-181b-1-3p↓, connected to 5 pathways (Figure S1); RAS related 2 (RRAS2), de-repressed by miR-3150b-3p↓, connected to 7 pathways (Figure S2); Heme oxygenase 1 (HMOX1), de-repressed by miR-873-5p↓ (validated, TarBase), connected to 3 pathways (Figure S3); and NFE2 like bZIP transcription factor 2 (NFE2L2), de-repressed by miR-144- 3p (validated by dual luciferase reporter assay with site-directed mutagenesis), connected to NRF2-mediated Oxidative Stress Response pathway (Sangokoya *et al*., 2010) (Figure S4).

### Stress granule pathway shows strongest inhibitory bias

Among the enriched pathways, Stress Granule Signalling showed the strongest inhibitory directionality in the miRNA network (z = −2·53, −log(P) = 2·75) and the fourth most perturbed in the full transcriptome (z = −4·32, −log(P) = 19·9; Figure 2A). Ten miRNAs converged on 11 target genes within this pathway (Figure 4). Eight upregulated miRNAs repressed 9 SG components, while two downregulated miRNAs de-repressed *EIF4E* and *PIK3R3*. The 8:2 inhibitory ratio was the strongest directional bias of any pathway in the network.

**Figure 4.**
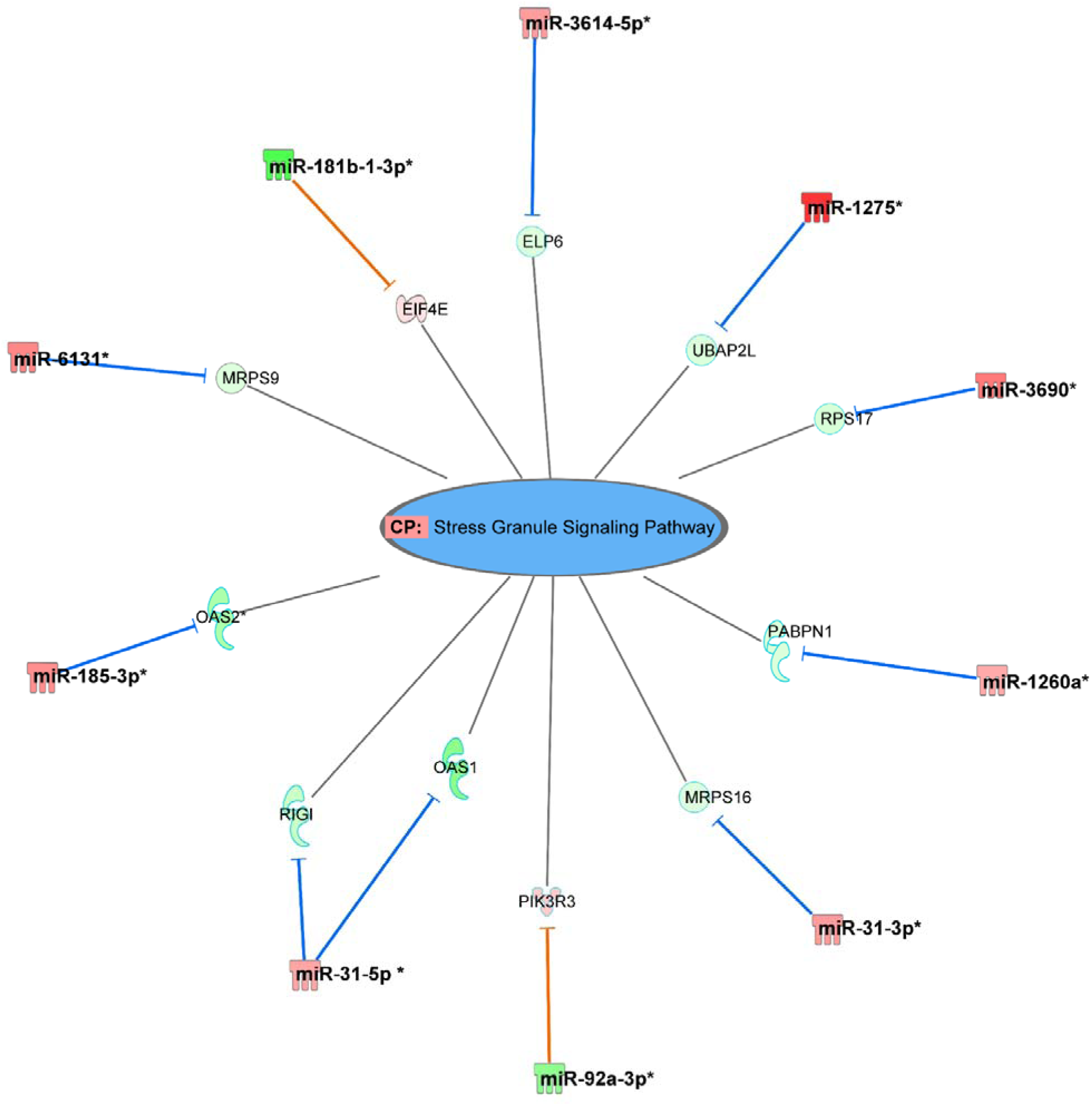
miRNA regulation of Stress Granule Signaling in heatstroke. IPA network overlay showing 10 miRNAs converging on 11 target genes (z = −2.53). The 8:2 inhibitory ratio produces net suppression despite concurrent ISR activation.

### mTOR ambivalence resolved by the miRNA layer

*mTOR* Signalling was highly enriched but functionally directionless in the full transcriptome (z = +0·19, −log(P) = 11·1). The miRNA network resolved this ambivalence to significant activation (z = +2·24; Figure 5). Eight miRNAs converged on 8 target genes with a 5:3 activating ratio: five downregulated miRNAs de-repressed signalling components (*PIK3R3, EIF4E, RRAS2, HMOX1, RHOB*), while three upregulated miRNAs repressed ribosomal proteins (*RPS17, MRPS16, MRPS9*).

**Figure 5.**
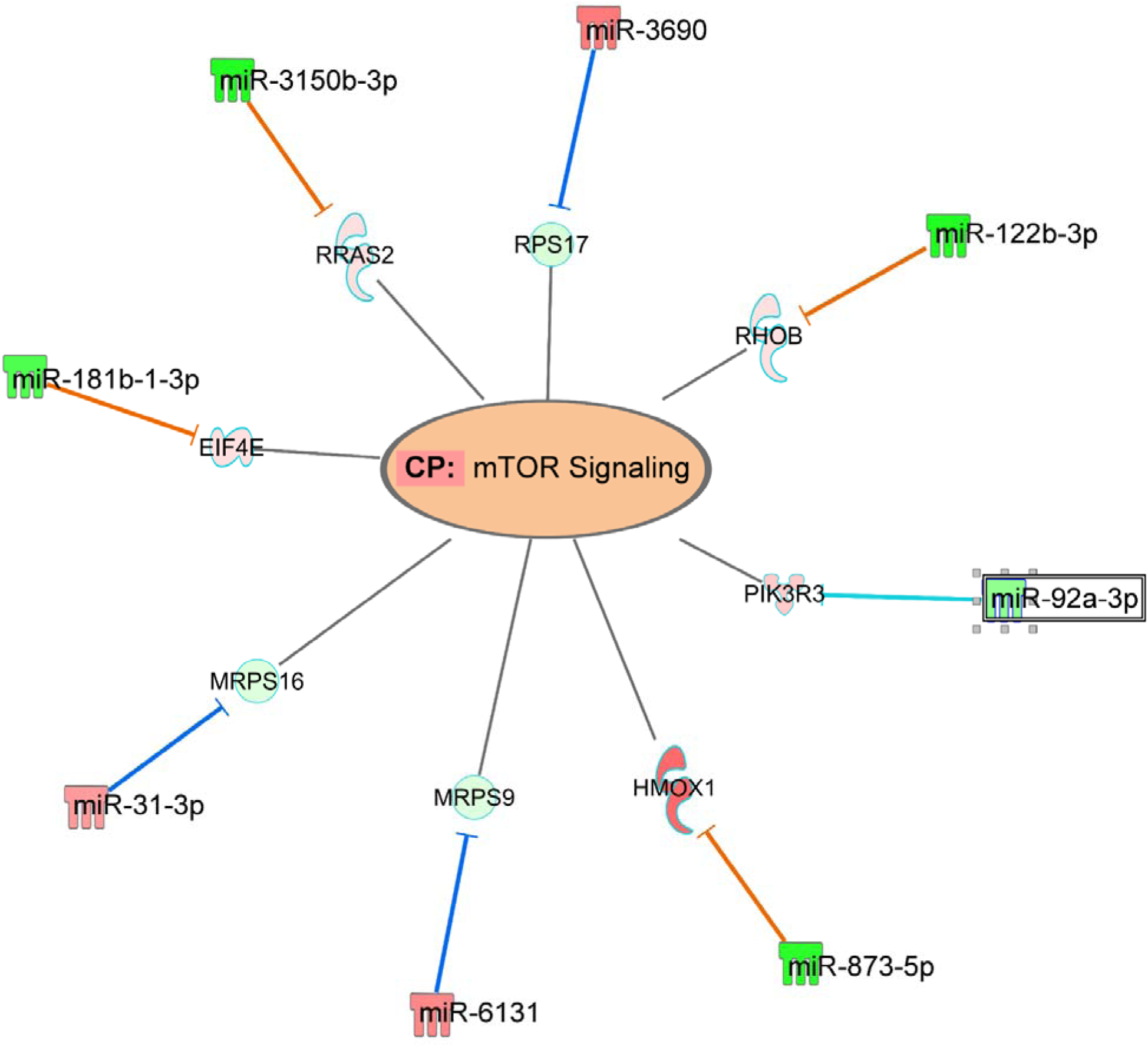
miRNA regulation of mTOR signaling in heatstroke. IPA network overlay showing 8 miRNAs converging on 8 target genes within mTOR Signaling (z = +2.24). Orange/cyan edges: de-repression by downregulated miRNAs (green nodes); blue edges: repression by upregulated miRNAs (red/pink nodes). Five downregulated miRNAs de-repress signaling components (*PIK3R3*, *EIF4E*, *RRAS2*, *HMOX1*, *RHOB*), while three upregulated miRNAs repress ribosomal proteins (*RPS17*, *MRPS16*, *MRPS9*). The 5:3 activating ratio resolves the transcriptomic ambivalence observed for mTOR in the full transcriptome (z = +0.19, Fig. 2A), producing predicted pathway activation through the miRNA regulatory layer.

### Fatty-acid β-oxidation attenuation and NRF2 activation

Three upregulated miRNAs targeting *HADH, ACAA2, THEM4* and *DBI* were associated with suppression of fatty-acid β-oxidation, with no detectable activating inputs in the miRNA-regulated pathway (Figure S6). Concurrently, insulin receptor signalling met activation criteria through six miRNAs with no inhibitory inputs, a pattern compatible with greater reliance on glucose-linked acute stress metabolism. NRF2 received input from eight miRNAs through six gene nodes, with seven activating and one inhibiting input, a redundancy unique among the 10 stringent pathways.

### Two opposing miRNA programmes define the regulatory architecture

The 26 miRNAs organised into two coherent, opposing programmes (Figure 6). The downregulated set acted through de-repression to activate stress-survival signalling, whereas the upregulated set acted through repression to suppress stress-granule and fatty-acid β-oxidation machinery. Five hub effectors carried most of the activating output: PIK3R3 (9 pathways), RRAS2 (7), EIF4E (5), HMOX1 (3), and NFE2L2 (NRF2 direct). Notably, five nodes PIK3R3, EIF4E, RPS17, MRPS16, and MRPS9 appeared in both activated and inhibited pathways, indicating that the two programmes were not independent but converged on a shared regulatory core.

**Figure 6.**
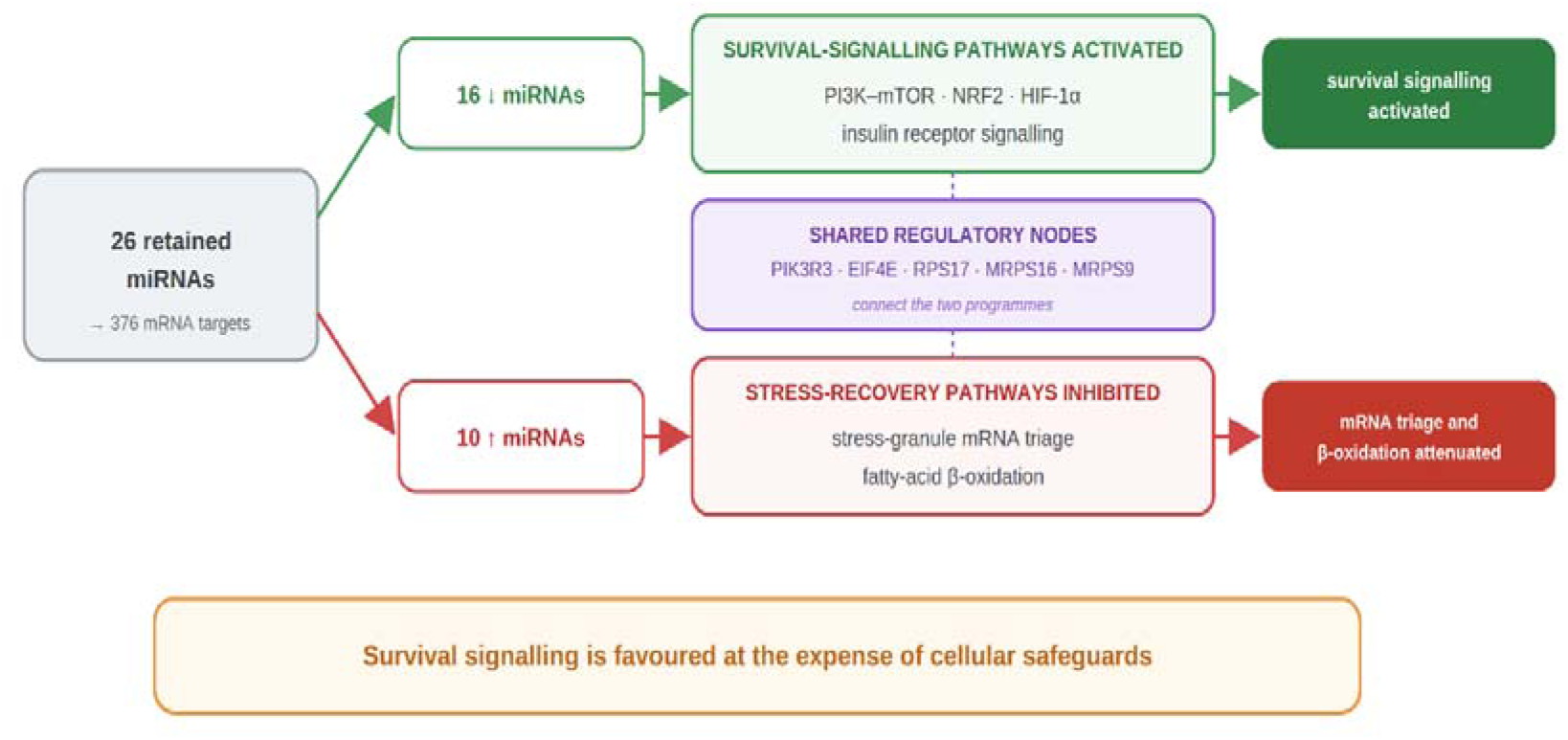
Two coherent regulatory programmes in heatstroke. Schematic summary of the two opposing miRNA-regulated programmes identified from the 26 retained miRNAs and 376 mRNA targets. Sixteen downregulated miRNAs were associated with activation of survival-signalling pathways, including PI3K–mTOR, NRF2, HIF-1α and insulin receptor signalling. Ten upregulated miRNAs were associated with inhibition of stress-recovery pathways, including stress-granule mRNA triage and fatty-acid β-oxidation. Shared regulatory nodes, including PIK3R3, EIF4E, RPS17, MRPS16 and MRPS9, connected the two programmes. These findings suggest a post-transcriptional trade-off in human heatstroke, in which short-term stress-survival signalling occurs alongside reduced protein quality control, metabolic flexibility and recovery capacity under extreme heat.

## Discussion

The present study extends our previous human heatstroke transcriptomic analysis by identifying a post-transcriptional miRNA layer that organises the acute PBMC response to severe heat injury. The principal physiological finding is a regulatory dissociation between activated stress-survival signalling and attenuated proteostasis and metabolic safeguards. Specifically, the miRNA-regulated network indicates suppression of stress granule signalling despite concurrent activation of ISR and UPR transcriptional programmes, suggesting uncoupling of processes that are normally tightly coordinated during cellular stress.

A key finding is an inferred uncoupling between ISR/UPR activation and the transcriptional programme supporting stress-granule assembly. In the canonical ISR, eIF2α phosphorylation triggers polysome disassembly, and released mRNP complexes nucleate into stress granules that function as mRNA triage centres. (Kedersha & Anderson, 2002; Anderson & Kedersha, 2008; Protter & Parker, 2016) In heatstroke, however, translational arrest extends beyond eIF2α-mediated suppression to structural downregulation of the translational machinery itself, yet the 8:2 inhibitory ratio in SG Signalling indicates that SG pathway components are suppressed rather than induced, despite concurrent ISR activation. This dissociation has experimental precedent: specific stresses, knockdown of SG nucleators and viral strategies that inhibit SG assembly. (Kedersha *et al*., 2013; Lu *et al*., 2021; Zheng *et al*., 2021)

We also found that reduced miR-181b-1-3p expression is consistent with de-repression of *EIF4E*, a key cap-binding factor required for canonical translation initiation.(Sonenberg & Hinnebusch, 2009) This creates a potential paradox: although *EIF4E* availability may increase, global translation remains strongly suppressed by ISR-mediated eIF2α phosphorylation. Elevated *EIF4E* activity may therefore compete with the selective translation programme imposed by the ISR, and, together with reduced stress-granule–mediated mRNA triage, favour poorly selective translation initiation under conditions of limited chaperone capacity and ATP, potentially increasing proteotoxic burden. (Wek *et al*., 2006; PakosCZebrucka *et al*., 2016).

Our data show that the suppression of fatty acid β-oxidation without detectable activating signals in pathway analysis is metabolically consistent with mitochondrial dysfunction, since β-oxidation in damaged mitochondria can generate toxic acylcarnitine intermediates.(Schooneman *et al*., 2013; Houten *et al*., 2016) The concurrent activation of insulin receptor signalling, without inhibitory inputs, is compatible with a shift toward greater reliance on glucose metabolism. In contrast, NRF2 signalling shows strong redundancy (seven activating versus one inhibitory input), suggesting that antioxidant defence remains robustly maintained even as other stress-response pathways diverge.

Previous studies have established the role of miR-92a-3p as a modulator of *PI3K/mTOR* signalling in cancer, where it targets *PTEN* to activate *PI3K/AKT/mTOR* pathways.(Lu *et al*., 2017; Yang *et al*., 2020; Wang *et al*., 2021) In ischemia, inhibition of miR-92a-3p enhances angiogenesis and functional recovery.(Bonauer *et al*., 2009) In heatstroke, the inferred regulatory effect appears to operate through de-repression of *PIK3R3* rather than *PTEN* or pharmacological inhibition, while converging on the same *PI3K*-dependent survival signalling. Heatstroke shares several pathophysiological features with ischaemia–reperfusion injury, including cardiovascular collapse, impaired tissue perfusion, and systemic inflammation.(Bouchama *et al*., 2022) Heat stress also directly stabilises HIF-1α through an HSP90-dependent mechanism independent of hypoxia.(Katschinski *et al*., 2002) The protective effects of miR-92a inhibition reported in ischemic tissues are therefore consistent with the *PI3K*-associated survival programmes observed in our analysis, suggesting that related regulatory mechanisms may operate across different forms of acute tissue stress.

The pathway classes we identified, energy metabolism, the unfolded protein response, and *PI3K-Akt* signalling, converge with the rat heatstroke model, where miRNA-regulated disruption of these same systems distinguished animals with organ injury from those that recovered.(Permenter *et al*., 2019) That the same three pathway axes are independently disrupted by miRNAs in both a controlled rat model and human heatstroke suggests a conserved mammalian stress architecture rather than a species- or context-specific response. Critically, all rats reached the same core temperature, yet only those with extensive miRNA dysregulation developed organ injury at 48 h, indicating that the extent of post-transcriptional regulatory disruption, rather than thermal dose alone, distinguished animals that recovered from those that did not.(Permenter *et al*., 2019)

Taken together, the two programmes are biologically coherent. The miRNA-regulated network suggests that activation of mTOR/EIF4E-associated survival signalling occurs together with suppression of stress-granule signalling, while PI3K–Akt-related signalling is accompanied by attenuation of fatty-acid β-oxidation in the setting of mitochondrial dysfunction. (Kedersha *et al*., 2013; Houten *et al*., 2016) However, this configuration carries inherent risk. Predicted mTOR activation would be expected to suppress autophagy, yet neither autophagy (z = +0.42) nor mitophagy (z = −0.54) reached activation criteria in our data. (Kim *et al*., 2011; Saxton & Sabatini, 2017) Damaged mitochondria may therefore continue generating reactive oxygen species, progressively eroding NRF2 reserve.(Youle & Narendra, 2011; Dinkova-Kostova & Abramov, 2015)

We also noted that protein ubiquitination, despite strong enrichment (−log(P) = 9·32), remained directionless (z = −0·12), indicating transcriptional conflict rather than coordinated proteasomal engagement. Programme A is associated with active defence signalling that may temporarily mask ongoing damage. In contrast, Programme B is characterised by reduced expression of proteostasis safeguards, including stress-granule triage, autophagic clearance, and mitophagic quality control, that would normally buffer proteotoxic accumulation. If NRF2 and heat-shock protein capacity are exceeded, no secondary defence is transcriptionally evident, and multiple protective mechanisms may fail concurrently rather than sequentially.

Whether these two expression groups represent a coordinated regulatory module or independent responses to distinct aspects of heat injury cannot be determined from the present cross-sectional data. This configuration provides a framework that may help explain the failure of a robust HSR to prevent severe heatstroke. (Bouchama *et al*., 2023). From a clinical perspective, the therapeutic window for cooling may depend not only on temperature but on the duration of this regulatory configuration, a hypothesis requiring prospective validation. Specific miRNAs such as miR-92a-3p may represent candidate biomarkers. The miRNA–pathway relationships suggest therapeutic levers: partial *mTOR* modulation to restore autophagic clearance, ISR agonists to restore SG formation, or targeted miRNA modulation, where antagomir inhibition of miR-92a has already demonstrated functional recovery in murine ischaemia models.(Bonauer *et al*., 2009; Hinkel *et al*., 2013; Saxton & Sabatini, 2017; Wu *et al*., 2024) These remain hypothesis-generating observations requiring validation in independent cohorts and longitudinal clinical studies.

Our study has several limitations. The analysis focuses on the admission time point before cooling therapy in 17 heatstroke patients and 16 controls for miRNA (19 and 19 for mRNA); miRNA trajectories during recovery and their relationship to clinical outcomes remain to be defined. miRNA–mRNA pairs were restricted to experimentally validated and high-confidence predicted interactions within the IPA miRNA Target Filter; a subset of these pairs have additional independent support in miRTarBase, including three validated by reporter assay.(Cui *et al*., 2025) Transcriptomic data reflect mRNA abundance, not protein levels or pathway flux. PBMCs were intentionally studied as circulating immune effector cells and as the clinically accessible compartment during emergency presentation; they are not proposed as surrogates for organ-specific injury, but as a window into the systemic post-transcriptional response to extreme heat. This design is therefore appropriate to the study objective.

## ADDITIONAL INFORMATION

### Data availability statement

De-identified microarray expression data are publicly available in the Gene Expression Omnibus (GEO: GSE317880; https://www.ncbi.nlm.nih.gov/geo/query/acc.cgi?acc=GSE317880). Small RNA sequencing data have also been deposited in GEO under accession number GSE324277 and will be released upon publication; reviewer access will be provided during peer review if required.

miRNA expression profiles and seed family annotations are provided in Supplementary Table S1. The complete IPA microRNA Target Filter-derived list of 414 inverse-expression miRNA–mRNA pairs, including the target-filter settings, is provided in Supplementary Table S2.

Individual participant clinical data cannot be shared because the study involved a vulnerable population (critically ill Hajj pilgrims), and the institutional review board approval restricts data sharing to variables reported in aggregate in the manuscript. Summary-level clinical data are provided in Table 1.

A data dictionary is not applicable because all variables are defined in the manuscript. No additional related documents (such as a study protocol or statistical analysis plan) are available.

Genomic datasets will be available indefinitely through GEO without restriction. Requests for collaboration involving additional analyses may be directed to the corresponding author.

### Author Contributions

AB was responsible for the study concept and design. SAM, MLA, and SSM collected clinical samples and data under supervision by AB. MG and RH conceived the analytical method and performed bioinformatic analysis. MA performed small RNA sequencing. MG, RH and AB performed miRNA–mRNA integration and pathway analysis. MG, SAM, MLA, SSM, SY and AB interpreted the data. MG and AB wrote the manuscript. Every author had full access to the dataset. MG and SAM directly accessed and verified the underlying data reported in the manuscript. All authors contributed to the interpretation of findings, provided critical revision of the manuscript for important intellectual content, and approved the final version for publication. AB had final responsibility for the decision to submit for publication.

## Acknowledgements

This work was supported by King Abdullah International Medical Research Center (grant RC18/373/R). We thank Haitham Alkadi and Deemah Alwadaani for microarray hybridisation and the Global Center for Mass Gathering Medicine, Ministry of Health, Riyadh, Saudi Arabia for logistical support during field sampling.

