## Supplemental Table and Figures for "MicroRNA regulation of stress-survival signalling and protein quality control in human heatstroke"

### Supplemental materials

Table S1. Complete miRNA regulatory hub analysis — 26 miRNAs with high-confidence inverse expression pairing to 376 mRNA targets.

| **Rank** | **miRNA** | **Seed** | **Targets** | **log₂FC** | **Direction** |
| --- | --- | --- | --- | --- | --- |
| 1 | mir-185-3p | GGGGCUG | 55 | +1.595 | ↑ UP |
| 2 | mir-1275 | UGGGGGA | 51 | +2.906 | ↑ UP |
| 3 | mir-92a-3p/mir-363-3p | AUUGCAC | 31 | -1.058 | ↓ DOWN |
| 4 | mir-1260a | UCCCACC | 29 | +1.197 | ↑ UP |
| 5 | mir-31-5p | GGCAAGA | 25 | +1.272 | ↑ UP |
| 6 | mir-181a-5p | ACAUUCA | 22 | -1.566 | ↓ DOWN |
| 7 | mir-3150b-3p | GAGGAGA | 20 | -2.588 | ↓ DOWN |
| 8 | mir-181b-1-3p | UCACUGA | 19 | -1.682 | ↓ DOWN |
| 9 | mir-92a-1-5p | GGUUGGG | 19 | +1.215 | ↑ UP |
| 10 | mir-4286 | CCCCACU | 18 | +1.955 | ↑ UP |
| 11 | mir-338-3p | CCAGCAU | 15 | -1.211 | ↓ DOWN |
| 12 | mir-181a-2-3p | CCACUGA | 13 | -1.671 | ↓ DOWN |
| 13 | mir-122b-3p | AACACCA | 13 | -2.633 | ↓ DOWN |
| 14 | mir-196a-5p | AGGUAGU | 11 | -1.953 | ↓ DOWN |
| 15 | mir-6131 | GCUGGUC | 11 | +1.695 | ↑ UP |
| 16 | mir-31-3p | GCUAUGC | 9 | +1.418 | ↑ UP |
| 17 | mir-144-3p | ACAGUAU | 8 | -1.563 | ↓ DOWN |
| 18 | mir-3614-5p | CACUUGG | 8 | +1.397 | ↑ UP |
| 19 | mir-3690 | CCUGGAC | 7 | +1.892 | ↑ UP |
| 20 | mir-10399-5p | AUUACAG | 6 | -1.068 | ↓ DOWN |
| 21 | mir-10a-5p | ACCCUGU | 6 | -2.133 | ↓ DOWN |
| 22 | mir-3611 | UGUGAAG | 4 | -1.073 | ↓ DOWN |
| 23 | mir-4422 | AAAGCAU | 4 | -2.025 | ↓ DOWN |
| 24 | mir-873-5p | CAGGAAC | 4 | -2.014 | ↓ DOWN |
| 25 | mir-4521 | CUAAGGA | 4 | -1.801 | ↓ DOWN |
| 26 | mir-451a | AACCGUU | 2 | -1.692 | ↓ DOWN |

**Figure S1. miR-181b-1-3p regulatory network and EIF4E multi-pathway node.**

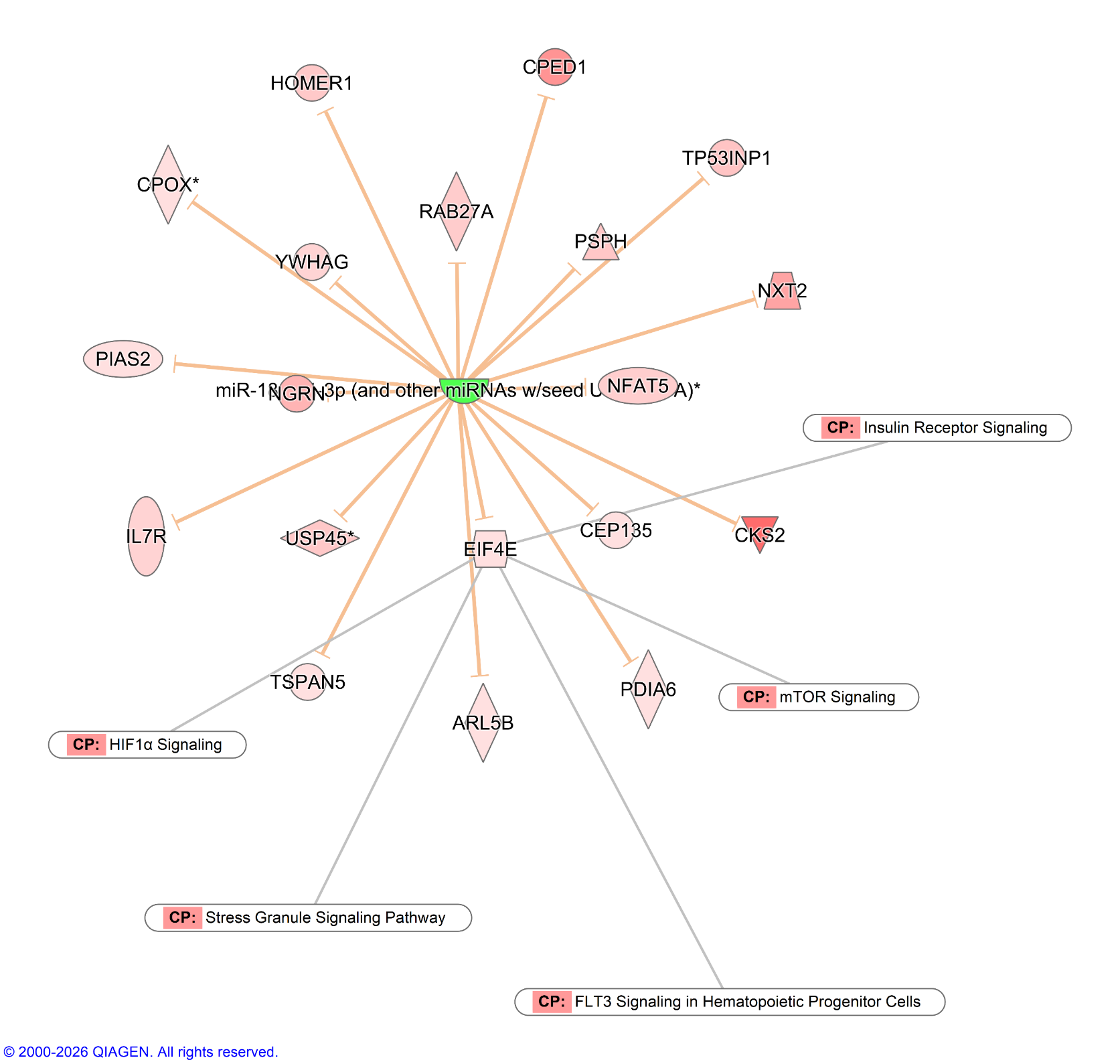

**Figure S2. miR-3150b-3p regulatory network and RRAS2 multi-pathway node.**

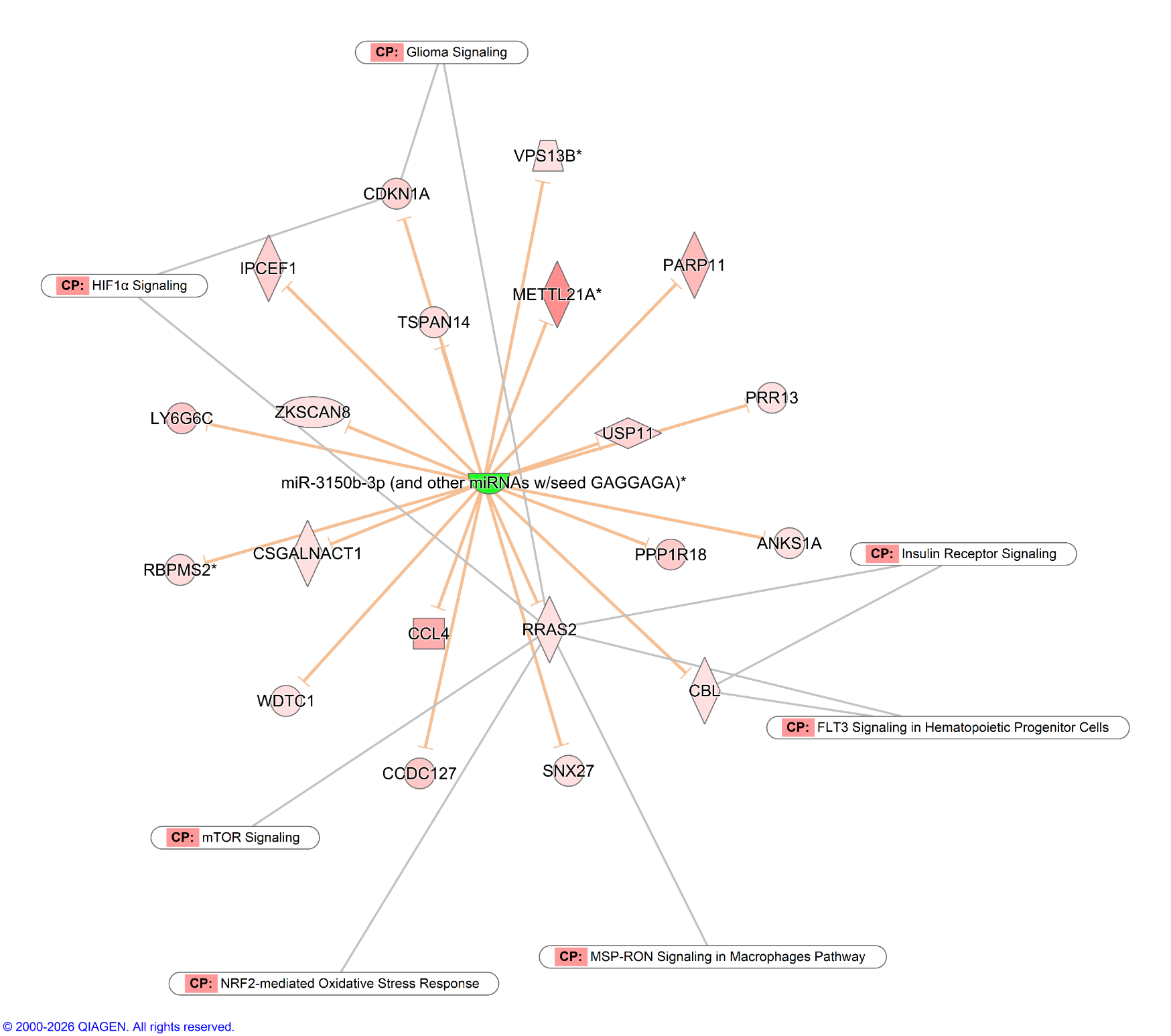

**Figure S3. miR-873-5p regulatory network and HMOX1 multi-pathway node.**

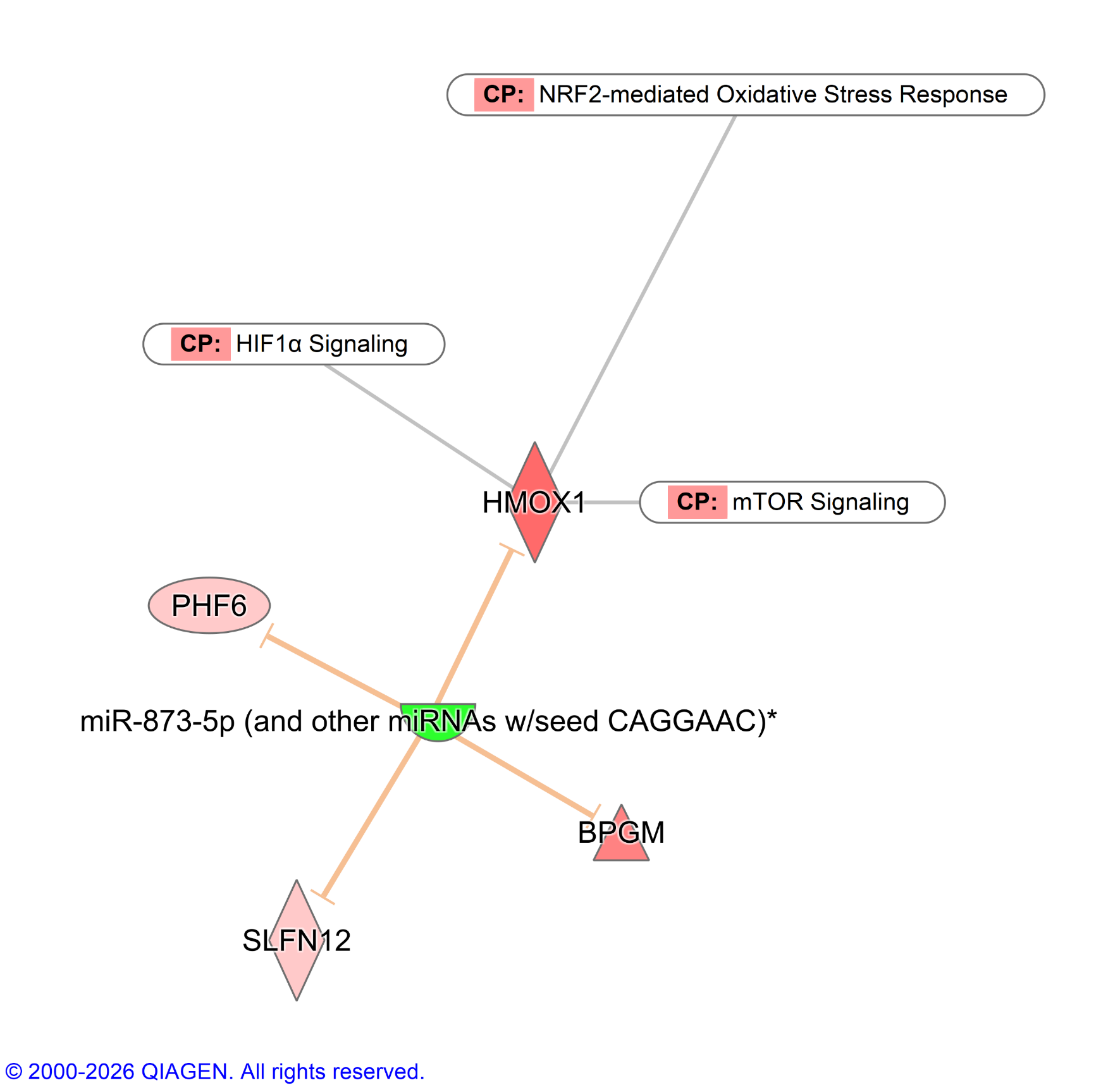

**Figure S4. miR-144-3p regulatory network and NFE2L2 pathway node.**

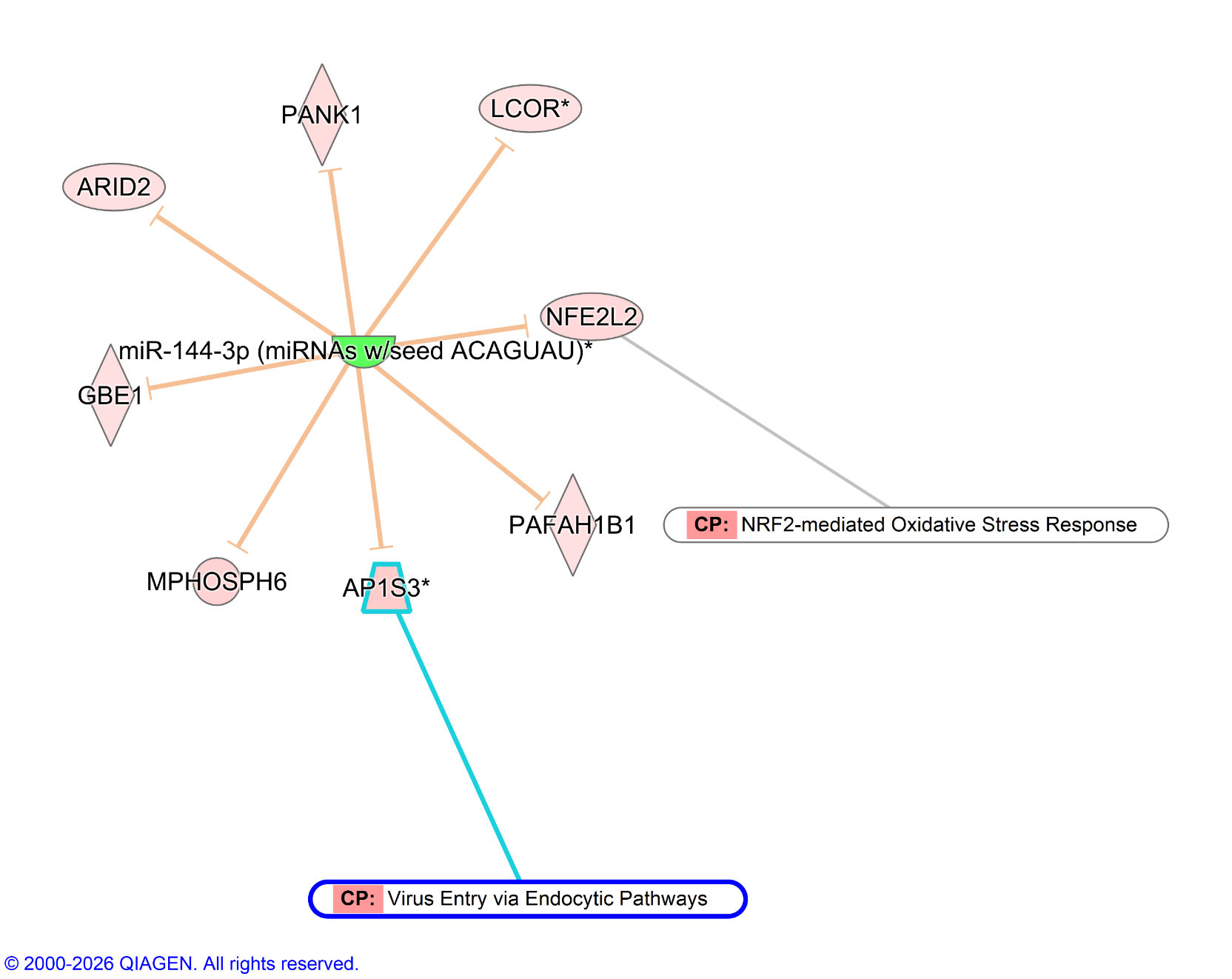

**Figure S5. miR-31-3p regulatory network and BACH1 repression.**

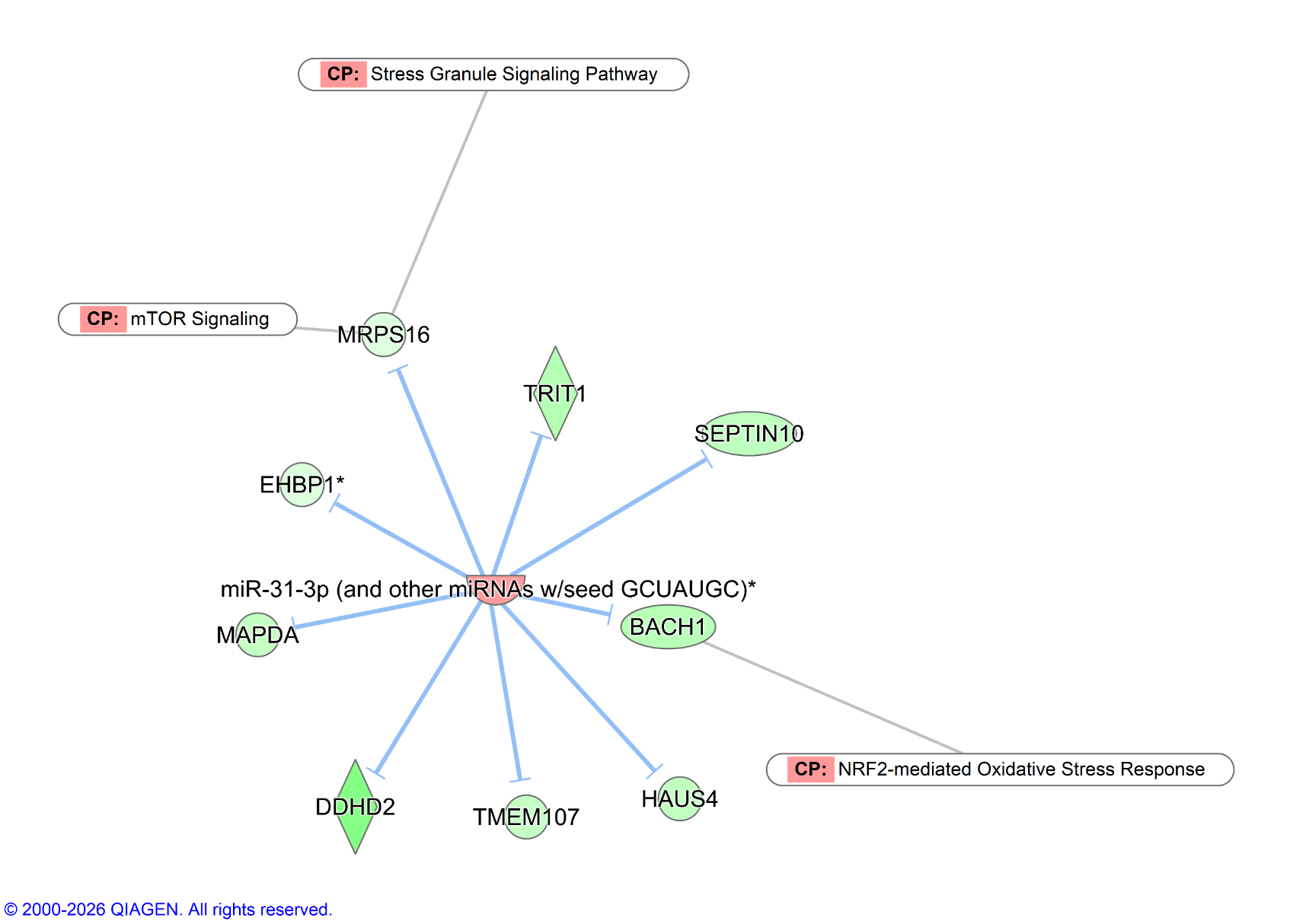

**Figure S6. miRNA regulation of Mitochondrial Fatty Acid β-Oxidation in heatstroke.**

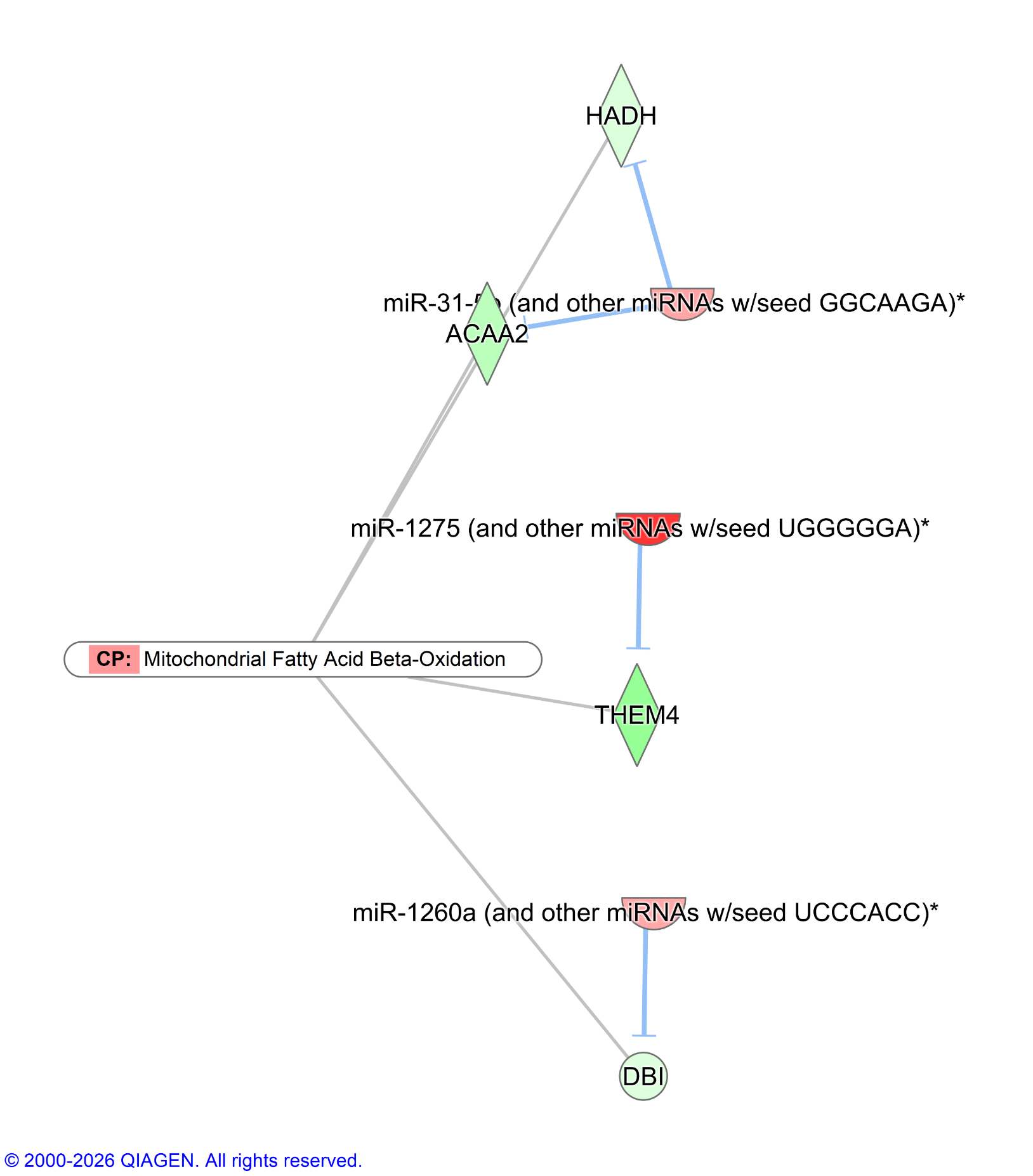

**Supplemental figure legends**

**Figure S1. miR-181b-1-3p regulatory network and EIF4E multi-pathway node.**

Network overlay of miR-181b-1-3p (downregulated) and its 19 de-repressed mRNA targets. Orange edges indicate de-repression (miRNA↓ → mRNA↑). EIF4E connects to 5 of 10 stringent canonical pathways (pink boxes): HIF1α Signaling, mTOR Signaling, FLT3 Signaling in Hematopoietic Progenitor Cells, Insulin Receptor Signaling, and Stress Granule Signaling Pathway. Gray lines indicate gene–pathway membership. EIF4E encodes the rate-limiting cap-binding protein for translation initiation, positioning it at the intersection of translational control and stress granule dynamics.

**Figure S2. miR-3150b-3p regulatory network and RRAS2 multi-pathway node.**

Network overlay of miR-3150b-3p (downregulated) and its 20 de-repressed mRNA targets. Orange edges indicate de-repression (miRNA↓ → mRNA↑). RRAS2 connects to 7 stringent canonical pathways: HIF1α, NRF2, mTOR, Insulin Receptor, FLT3, Glioma Invasiveness, and MSP-RON Signaling. Gray lines indicate gene–pathway membership

**Figure S3. miR-873-5p regulatory network and HMOX1 multi-pathway node.**

Network overlay of miR-873-5p (downregulated) and its 4 de-repressed mRNA targets. Orange edges indicate de-repression (miRNA↓ → mRNA↑). HMOX1 connects to 3 stringent canonical pathways (pink boxes): NRF2-mediated Oxidative Stress Response, HIF1α Signaling, and mTOR Signaling. HMOX1 encodes heme oxygenase-1, a cytoprotective enzyme and validated target of miR-873-5p (TarBase).

**Figure S4. miR-144-3p regulatory network and NFE2L2 pathway node.**

Network overlay of miR-144-3p (downregulated) and its 8 de-repressed mRNA targets. Orange edges indicate de-repression (miRNA↓ → mRNA↑). NFE2L2 (Nrf2) connects to NRF2-mediated Oxidative Stress Response (gray line). AP1S3 connects to Virus Entry via Endocytic Pathways (cyan line). NFE2L2 encodes the master transcription factor for antioxidant response element–driven gene expression.

**Figure S5. miR-31-3p regulatory network and BACH1 repression.**

Network overlay of miR-31-3p (upregulated) and its 9 repressed mRNA targets. Blue edges indicate repression (miRNA↑ → mRNA↓). BACH1 connects to NRF2-mediated Oxidative Stress Response. MRPS16 connects to Stress Granule Signaling and mTOR Signaling pathways.

**Figure S6. miRNA regulation of Mitochondrial Fatty Acid β-Oxidation in heatstroke.**

IPA network overlay showing 3 upregulated miRNAs converging on 4 target genes within Mitochondrial Fatty Acid β-Oxidation (z = −2.00). Blue edges: repression by upregulated miRNAs (red nodes). miR-31-5p represses both HADH and ACAA2, miR-1275 represses THEM4, and miR-1260a represses DBI. All inputs are inhibitory (3:0 ratio), representing an exclusively inhibitory signal within the network. Gray lines indicate gene–pathway membership.
